# mTORC2 contributes to murine systemic autoimmunity

**DOI:** 10.1101/2021.03.27.437347

**Authors:** Xian Zhou, Haiyu Qi, Meilu Li, Yanfeng Li, Xingxing Zhu, Shreyasee Amin, Mariam Alexander, Catherine Diadhiou, Anne Davidson, Hu Zeng

**Affiliations:** Division of Rheumatology, Department of Medicine, Mayo Clinic Rochester, MN 55905, USA; Department of Rheumatology, Beijing Friendship Hospital, Capital Medical University, Beijing, 100050, P. R. China; Department of Dermatology, the Second Hospital of Harbin Medical University, Harbin Medical University, Harbin, 150001, P. R. China; Division of Laboratory Medicine and Pathology, Mayo Clinic Rochester, MN 55905, USA; Institute of Molecular Medicine, Feinstein Institutes for Medical Research, Manhasset, NY 11030, USA; Donald and Barbara Zucker School of Medicine at Hofstra/Northwell, Manhasset, NY 11030, USA; Department of Immunology, Mayo Clinic Rochester, MN 55905, USA

## Abstract

The development of many systemic autoimmune diseases, including systemic lupus erythematosus, is associated with overactivation of the type I interferon (IFN) pathway, lymphopenia, and increased follicular helper T (Tfh) cell differentiation. However, the cellular and molecular mechanisms underlying these immunological perturbations remain incompletely understood. Here we show that the mechanistic target of rapamycin complex 2 (mTORC2) promotes Tfh differentiation and disrupts Treg homeostasis. Inactivation of mTORC2 in total T cells, but not in Tregs, greatly ameliorated the immunopathology in a systemic autoimmunity mouse model. This was associated with reduced Tfh differentiation, B cell activation, and reduced T cell glucose metabolism. Finally, we show that type I IFN can synergize with TCR ligation to activate mTORC2 in T cells, which partially contributes to T cell lymphopenia. These data indicate that mTORC2 may act as downstream of type I IFN, TCR, and costimulatory receptor ICOS, to promote glucose metabolism, Tfh differentiation, and T cell lymphopenia, but not to suppress Treg function in systemic autoimmunity. Our results suggest that mTORC2 might be a rational target for systemic autoimmunity treatment.

## INTRODUCTION

The cellular and molecular mechanisms leading to systemic autoimmunity remain poorly understood. A prototype of systemic autoimmunity is systemic lupus erythematosus (SLE). Two major immunological alterations have been identified in lupus, overactivation of the type I IFN signaling pathway and a loss of balance between effector T cells, particularly follicular helper T (Tfh) cells, and regulatory CD4^+^ T cells (Tregs)^1–3^. Therapeutical options like blocking type I IFN, or its receptor have shown clinical efficacy^4^. Yet, the specific cellular and molecular mechanisms by which type I IFN promotes SLE pathology remain incompletely understood. Recent studies using murine lupus models have demonstrated that type I IFN signaling in CD4^+^ T cells are required for development of lupus pathology^5,6^. In addition, it is well established that type I IFN can block T cell egress from lymph nodes and induce lymphopenia partly by upregulating the expression of CD69^7,8^, a marker that is also associated with lupus pathology^9,10^. Clinical studies^11^ and large-scale single cell RNA sequencing analysis^12^ confirmed the inverse association between type I IFN activity and the abundance of circulating lymphocytes or naïve CD4^+^ T cells in SLE patients. However, the type I IFN signaling pathway in CD4^+^ T cells remains incompletely understood and it is unclear how type I IFN overactivation may be linked to T cell lymphopenia in lupus.

Increased differentiation of Tfh cells has been associated with SLE development and severity in mouse models and human patients^13,14^. Germinal center (GC) Tfh cells, distinguished by expression of the transcription factor BCL6, chemokine receptor CXCR5, and immune checkpoint receptor PD-1, are a specialized effector T cell (Teff) lineage that stimulates B cells in GC to undergo affinity maturation and clonal expansion. The costimulatory receptor ICOS delivers critical signals for Tfh differentiation^15^. In addition, a special subset of Tfh cells with low expression of CXCR5 and P-selectin ligand, PSGL-1, are localized in extrafollicular (EF) to promote generation of plasmablasts, which produce antibodies with modest affinity^16–18^. Both GC and EF responses may contribute to SLE development^19^. In contrast, impaired Treg function is observed in some SLE patients^20^. However, what drives the imbalance between Tfh and Tregs in lupus is not well understood.

Mechanistic target of rapamycin (mTOR) is a central metabolic pathway, which consists of two complexes, mTORC1 and mTORC2, with scaffolding molecules RAPTOR (encoded by *RPTOR*) and RICTOR (encoded by *RICTOR*) as their defining components, respectively^21^. mTORC1 is critical for naïve T cell quiescence exit and proliferation, and all effector T cells differentiation^22^. In contrast, mTORC2 is essential for Tfh differentiation, but dispensable for T cell activation and modestly contributes to other effector T cell lineages^22,23^. Ligation of CD3 and ICOS activates mTORC2, which in turn promotes glucose metabolism in Tfh cells^24^. In terms of Tregs, mTORC1 maintains natural or thymic derived Treg (nTreg, or tTreg) functional competency^25–28^. Overactivation mTORC2 impairs Treg suppressive function and stability, especially their ability to suppress Tfh cells, which leads to systemic autoimmune disease with lupus characteristics^29,30^. Inhibition of mTORC2 in Tregs can partly restore immune tolerance in the absence of mTORC1^25^, or FOXP3^31^. Therefore, mTORC2 signaling controls the balance between Tfh and Treg by promoting the former and suppressing the latter. These fundamental discoveries suggest that targeting mTORC2 in T cells might benefit lupus disease.

Several studies have examined the contribution of mTORC1 to SLE development^32^. How mTORC2 contributes to systemic autoimmunity has been much less understood. A significant increase of phosphorylation of AKT at serine 473, an established direct target of mTORC2, was observed in T cells from lupus patients and it was associated with disease severity^33,34^. These studies suggest that mTORC2 overactivation might be a potential contributing factor for lupus. The underlying mechanisms, however, remain obscure. Here, we tested the hypothesis that genetic targeting mTORC2 in T cells may improve systemic autoimmunity in *C57BL/6-Fas^Lpr^* (Lpr) mice. We show that Lpr T cells have an elevated baseline activity of mTORC2. Loss of *Rictor* specifically in T cells significantly improves immunopathology in Lpr mice, including T cell lymphopenia and generation of autoantibodies, which is associated with normalization of the balance between Tfh cells and Tregs. Surprisingly, mTORC2 contributes to systemic autoimmunity primarily through promoting Tfh cells, but not suppressing Tregs, because specific deletion of *Rictor* in Tregs largely fails to ameliorate the immune activation in Lpr mice. Lastly, we demonstrate that type I IFN can synergize with TCR signaling to sustain mTORC2 activation and promote CD69 expression, which suppress T cell egress and lead to lymphopenia, partly in an mTORC2 dependent manner. Our data supports the notion that mTORC2 dependent Tfh expansion, but not Treg impairment, may contribute to the lupus-associated systemic immunopathology.

## MATERIAL and METHODS

### Mice

*Cd4*^Cre^*Rictor*^fl/fl^ mice have been described before^24^. *C57BL/6-Fas^Lpr^* and Foxp3^Cre^ mice were purchased from The Jackson Laboratory. Mice were bred and maintained under specific pathogen-free conditions in the Department of Comparative Medicine of Mayo Clinic with IACUC approval.

### ELISA

For autoantibodies detection in sera, 96-well plates (2596; Costar) were coated with ssDNA and chromatin (Sigma-Aldrich) in PBS overnight at 4°C used at a concentration of 1.6 μg/ml and 5 μg/ml, respectively. Plates were washed twice (0.05% Tween 20 in PBS), blocked with 5% blocking protein (Bio-Rad) for 1 h, and washed twice, and serially diluted sera samples were added for 1.5 h at 37 °C. Plates were washed four times and horseradish peroxidase (HRP)-conjugated detection Abs for IgG (Bethyl Laboratories) were added for 1 h at RT, washed four times, and tetramethylbenzidine (TMB) substrate was added. Reaction was stopped using 2N H2SO4 and read at 450 nm. The titers of mouse IgM, IgG, IgG2b and IgG3 (Fig. 1C) in sera were measured with the ELISA kits from Invitrogen.

**Figure 1.**
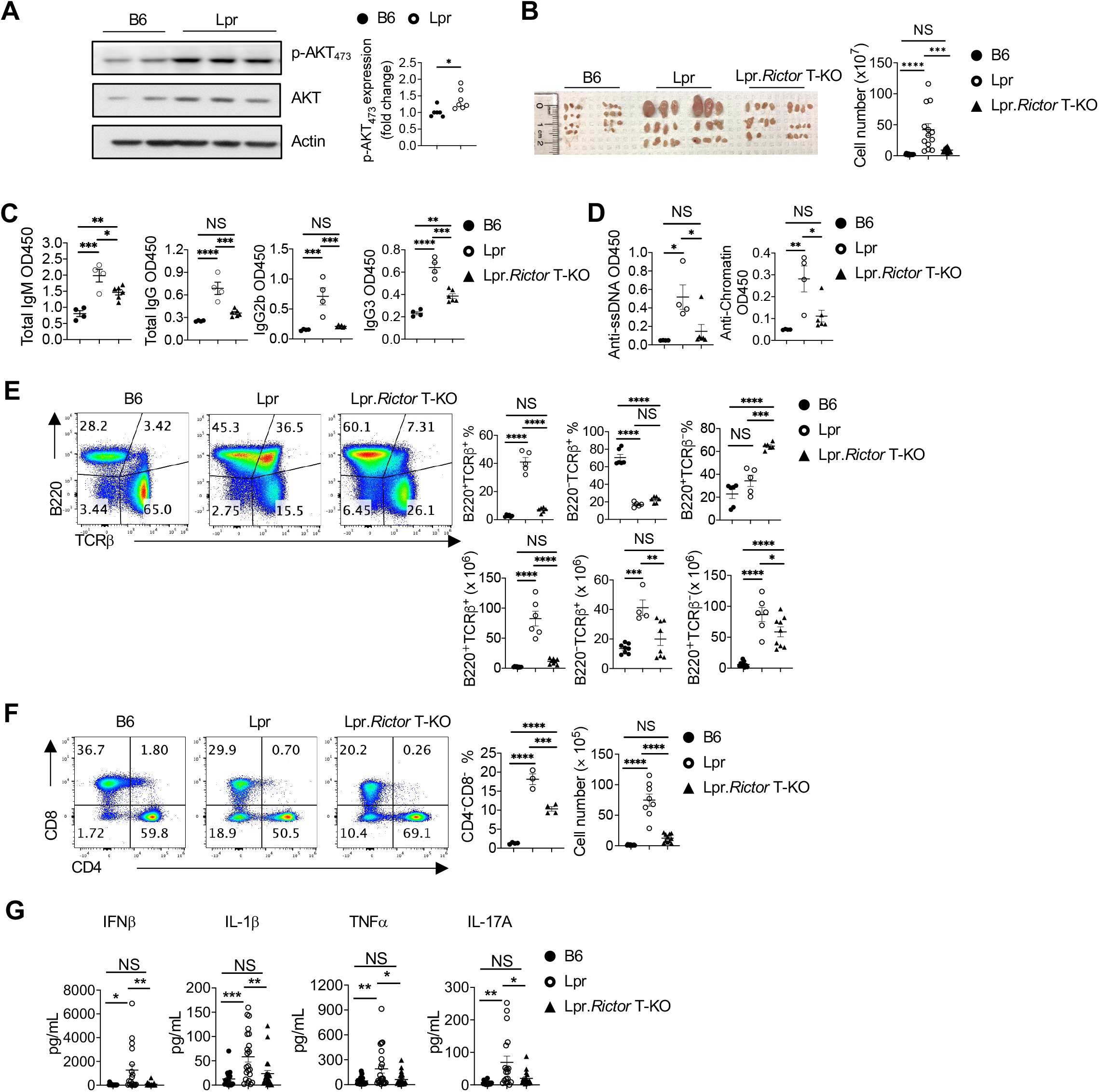
*Rictor* deletion in T cells rectifies immunopathology in Lpr mice. (A) Immunoblot analysis of p-AKT_473_ in B6 and Lpr CD4^+^ T cells isolated from peripheral lymph nodes (pLN). Right, summary of the relative p-AKT_473_ expression (normalized to that in B6 CD4^+^ T cells). (B) Image of peripheral lymph nodes taken from 6 months old B6, Lpr and Lpr.*Rictor* T-KO mice. Right, summary of total cellularity of lymph nodes. (C) Titers of total immunoglobulin (Ig) M, IgG, IgG2b and IgG3 were measured by ELISA. (D) Titers of anti-ssDNA (left) and anti-chromatin (right) were measured by ELISA. Samples were from 9 months old B6, Lpr and Lpr.*Rictor* T-KO mice. (E) Expression of B220 and TCRβ on lymphocytes. Right, the frequencies (upper panels) and absolute numbers (lower panels) of B220^+^TCRβ^+^, B220^-^TCRβ^+^, B220^+^TCRβ^-^ cells. (F) Flow cytometry analysis of CD4 and CD8 T cells among B220^-^TCRβ^+^ cells. Right, the frequency (left) and absolute number (right) of CD4^-^CD8^-^T cell population. (E) and (F) Cells were from peripheral lymph nodes (pLN) of 6 months old B6, Lpr and Lpr.*Rictor* T-KO mice. (G) The inflammatory cytokine levels in mouse sera samples collected from 5-6 months old B6, Lpr and Lpr.*Rictor* T-KO mice. NS, not significant; * *P* < 0.05, ** *P* < 0.01, *** *P* < 0.001, **** *P* < 0.0001. *p* values were calculated with one-way ANOVA. Results were presentative of 3 (A) independent experiment and pooled from at least 3 (A-G) independent experiments. Error bars represent SEM.

### Mouse immunoglobulin isotyping panel detection

The concentration of mouse immunoglobulin isotypes IgG1, IgG2a, and IgG2b (Fig. 4G) in sera were measured with the LEGENDplex mouse immunoglobulin isotyping panel, according to manufacturer’s instructions (Biolegend; cat# 740493).

### Immunofluorescence staining

Kidneys were obtained from 6-month-old B6, Lpr and Lpr. *Rictor*^fl/fl^ mice, and fixed in 4% paraformaldehyde (PFA) overnight, osmotically dehydrated in 20 % (w/v) sucrose, embedded in OCT, and cryosectioned to 6 μM. Sections were fixed in −20°C acetone for 2 min and blocked in PBS containing 8% goat serum, 1% bovine serum albumin (BSA) and 0.1% Tween 20 for 45 min. Deposition of immunoglobulin (IgG) and complement component 3 (C3) in the kidney tissue were stained with PE labeled goat anti-mouse IgG (Invitrogen) and FITC labeled rat anti-mouse C3 (Cedarlane) antibodies. Similarly, spleens were obtained from 6 months old mice, and processed as above. Frozen sections were fixed and blocked as described above, and incubated with primary antibodies Biotin Peanut Agglutinin (PNA) (Vectorlabs), purified rat anti-mouse CD3 (Biolegend) and PE labeled anti-mouse IgD (BD), or stained with Biotin anti-mouse CD138 (Biolegend) and PE labeled anti-mouse IgD at 4°C for 4 hours. Slides were then washed before streptavidin-Alexa Fluor 488 conjugate (ThermoFisher) and Alexa Fluor 647 labeled goat anti-rat secondary antibody (ThermoFisher) were added for 1 h at room temperature. Sections were covered with coverslip and visualized under an Olympus BX51 fluorescence microscope.

### Flow cytometry

Single-cell suspension was prepared by passing spleen and peripheral lymph nodes through a 70-μm nylon mesh. For analysis of surface markers, cells were stained in PBS containing 1% (w/v) BSA on ice for 30 min, with APC-labeled anti-ICOS (clone: 7E.17G9; BD), PE-Cy7-labeled anti PD-1 (clone: RMPI-30; BioLegend), Super Bright 600–labeled anti-CD4 (Clone: SK-3; eBioscience), BV510-labeld anti-CD8a (clone:53-6.7; BioLegend); APC-Cy7-labeled TCRβ (clone: H57-597; BioLegend), APC anti-CD162 (PSGL1; clone: 2PH1; BD), BV605-labeled anti-CD25 (clone: PC61; BioLegend), PE-Cy7–labeled anti-CD19 (clone: 6D5; BioLegend), BV605-labeled B220 (clone: RA3-6B2; BioLegend), PE-labeled anti-Fas (clone: Jo2; BD), BV785-labeled anti-CD138 (clone: 281-2; BioLegend), APC-labeled anti-IgD (clone: IA6-2; BioLegend), PerCP-Cyanine5.5-labeled anti-CD38 (clone: 90; BioLegend), FITC-labeled anti-IgG1 (clone: RMG1-1; BioLegend), APC-labeled anti-IL7Rα (clone: SB/199; BioLegend), FITC-labeled anti-CD69 (clone: H1.2F3; Tonbo Bio), PE-labeled anti-CD153 (clone: RM153; eBioscience), PE-labeled anti-CD73 (clone: TY/11.8; BioLegend) and APC-labeled anti-FR4 (clone: 12A5; BD). Cell viability was assessed using the Fixable Dye Ghost 540 (Tonbo Bioscience), or 7AAD. CXCR5 and PNA were stained with biotinylated anti-CXCR5 (clone: 2G8, BD Biosciences) or biotinylated peanut agglutinin (FL10-71; Vectorlabs), followed by staining with streptavidin-conjugated PE (BD Biosciences). Splenocytes and peripheral lymph node cells were cultured for 4 h at 37°C in complete medium (1640 RPMI + 10% FBS) containing ionomycin (1 μg/ml), PMA (50 ng/ml) and Monensin solution (1,000×; BioLegend) for cytokines detection. Intracellular cytokine staining of IFN-γ (clone: XMG1.2; eBioscience), IL-2 (clone: JES6-5H4; BioLegend) and IL-17A (clone: TC11-18H10.1; BioLegend) were performed using the BD Cytofix/Cytoperm and Perm/Wash buffers or, for transcriptional factors FoxP3 (clone: FJK-16s; eBioscience), Bcl6 (clone: K112-91, BD) and Ki67 (clone: SolA15; eBioscience) staining using the eBioscience Transcription Factor Staining Buffer Set. Phosflow staining for AF647 conjugated phosphor-AKT (S473) (D9E; Cell signaling) was performed using Phosflow Fix/Perm kit (BD Biosciences). Flow cytometry was performed on the Attune NxT (ThermoFisher) cytometer, and analysis was performed using FlowJo software (Tree Star).

### Immunoblotting

For immunoblotting, murine CD4^+^ T cells were isolated from mouse spleen and peripheral lymph node single cell suspension, and human CD4^+^ T cells were enriched from healthy donor PBMC with STEMCELL mouse or human CD4^+^ T cell isolation kits, respectively. Cells were lysed in lysis buffer with protease and phosphatase inhibitors (Sigma-Aldrich). Protein concentration in samples was quantified by BCA assay (Thermo Fisher Scientific) before loading the samples for electrophoresis and membrane transfer. The transferred membrane was blocked with TBST (0.1% Tween 20) containing 5% BSA for 1 h at room temperature. The membrane was incubated with primary antibodies overnight including anti-phospho-AKT (Ser473) (clone: D9E; Cell Signaling), AKT (pan) (Clone: 40D4; Cell Signaling), phospho-STAT1 (Tyr701) (clone: D4A7; Cell Signaling), phospho-STAT2 (Tyr690) (Cell Signaling), phospho-S6 (Ser235/236) (clone: D57.2.2E; Cell Signaling) and anti-β-actin (clone: 13E5; Sigma-Aldrich). Then, the membrane was washed and incubated with the corresponding secondary antibody for subsequent enhanced chemiluminescence (ECL; Thermo Fisher) exposure. The band intensity of all the immunoblot was analyzed by ImageJ software.

### Inflammatory cytokines detection

Murine sera were collected from 6-month-old B6, Lpr and Lpr.*Rictor* T-KO mice. The levels of inflammatory cytokines in serum were measured with the LEGENDplex Multi-Analyte Flow Assay Kit, according to manufacturer’s instructions (Biolegend, Mouse Inflammation Panel, cat# 740446).

### ICOS stimulation

Cell isolation and flow cytometry of lymphocytes were as described. One million CD4^+^ T cells were activated with plate coated anti-CD3 and anti-CD28 (Bio X Cell). After 2 days activation, T cells were removed from stimulation and rested overnight. Live cells were purified using lymphocyte isolation buffer (MP Biomedicals), followed by stimulation with plate bound anti-CD3 (2 μg/ml) and anti-ICOS (5 μg/ml; C398.4A; Biolegend) for 24 hours.

### Metabolic assay

The bioenergetic activities of the OCR and ECAR were measured using a Seahorse XFe96 Extracellular Flux Analyzed following established protocols (Agilent). Briefly, equal number of live CD4^+^ T cells were seeded at 200,000 cells/well on Cell-Tak (Corning) coated XFe96 plate with fresh XF media (Seahorse XF RPMI medium containing 10 mM glucose, 2 mM L-glutamine, and 1 mM sodium pyruvate, PH 7.4; all reagents from Agilent). Basal OCR and ECAR were measured in the presence of Oligomycin (1.5 μM; Sigma-Aldrich), FCCP (1.5 μM; Sigma-Aldrich), and Rotenone (1 μM; Sigma-Aldrich)/ Antimycin A (1 μM; Sigma-Aldrich) in Mito Stress assay. For ECAR measurement in ICOS secondary stimulated CD4^+^ T cells, glycolytic rate assay was performed according to the manufacturer’s manual (Agilent) in presence of Rotenone/Antimycin A and 2-DG (50 mM, Sigma-Aldrich)

### Poly(I:C) treatment and cell counting in peripheral blood

*Cd4*^Cre^*Rictor*^fl/fl^ and wild-type mice were given intraperitoneal injection of 200 μg poly(I:C) (Sigma-Aldrich) in 200 μL PBS, and blood lymphocyte counts were assessed 16 and 40 hours later. Briefly, to determine absolute blood cell counts, 50 μL peripheral blood with 5 μL EDTA was collected, lysed with 1 mL ACK lysing buffer (ThermoFisher), and washed with PBS. Samples were stained with anti-mouse CD19, CD4, CD8 and CD69 antibodies, washed with PBS once, suspended with 500 μL PBS and loaded to Attune NxT cytometer with the accurate acquisition of 250 μL samples. Cell counts were calculated per 5 μL blood.

### Statistical Analysis

Statistical analysis was performed using GraphPad Prism (version 8). P values were calculated with Student’s *t* test, or one-way ANOVA. *P* < 0.05 was considered significant.

## RESULTS

### Loss of mTORC2 in T cells improves immunopathology in murine systemic autoimmunity

Lpr mice recapitulate many of the systemic immunoproliferative phenotypes in human lupus disease without developing overt nephritis phenotypes^35^. We tested whether mTORC2 activity is altered in T cells from Lpr mice. Immunoblot analysis showed that CD4^+^ T cells from Lpr mice had elevated phosphorylation of AKT at Serine 473 (p-AKT_473_), a direct target of mTORC2, indicating overactivation of mTORC2 in Lpr T cells at baseline (Fig. 1A). To test the hypothesis that targeting mTORC2 may improve systemic autoimmunity, we crossed *Cd4*^Cre^*Rictor*^fl/fl^ mice, in which *Rictor* is specifically deleted and hence abrogates mTORC2 activity in T cells^24,25^, with Lpr mice to generate C57BL/6.*Fas^Lpr^Cd4*^Cre^*Rictor*^fl/fl^ (Lpr.*Rictor* T-KO) mice. RICTOR deficiency in T cells alone does not significantly impact immune cell homeostasis and overall mouse health^25,36,37^. Therefore, we focused on the comparison among B6, Lpr and Lpr.*Rictor* T-KO mice. *Fas^Lpr^* mutation drives dramatic lymphoid hyperplasia, including lymphadenopathy. We observed that deletion of *Rictor* in T cells significantly rescued lymphadenopathy phenotypes (Fig. 1B). Consistent with these findings, Lpr mice had increased total IgM and IgG, and isotypes IgG2b and IgG3 immunoglobulin levels (Fig. 1C) and increased anti-ssDNA and anti-chromatin autoantibodies (Fig. 1D), most of which were significantly reduced in Lpr.*Rictor* T-KO mice. Furthermore, we examined IgG and complement deposition in kidneys, which reflects renal involvement in autoimmunity. Lpr mice on B6 background have very mild kidney pathology compared to those on MRL/MpJ background^38^. Consistent with this observation, we observed a mild increase of IgG and complement C3 deposition in kidneys from Lpr mice. Deletion of *Rictor* reduced IgG and C3 deposition in Lpr.*Rictor* T-KO kidney, although the latter did not reach statistical significance (Supplemental Fig. 1A). Thus, inactivation of mTORC2 in T cells substantially attenuates lymphoid hyperplasia, antibody secretion, and renal immune complex deposition in a mouse model of systemic autoimmunity.

Consistent with the attenuated lymphoid hyperplasia in Lpr.*Rictor* T-KO mice, the accumulation of the aberrant B220^+^TCRβ^+^ cells in Lpr mice was nearly completely reversed in Lpr.*Rictor* T-KO mice (Fig. 1E). Although Lpr.*Rictor* T-KO mice had a significant increase of B220^+^TCRβ^-^ B cell percentage, the absolute number of this population modestly, but significantly, reduced in Lpr.*Rictor* T-KO mice compared to Lpr mice (Fig. 1E). B220^-^TCRβ^+^ T cell percentages were highly reduced in both Lpr and Lpr.*Rictor* T-KO mice, but Lpr mice had an increase of B220^-^ TCRβ^+^ T cell number likely due to the lymphoid hyperplasia, which was also reversed in Lpr.*Rictor* T-KO mice (Fig. 1E). Moreover, the accumulation of TCRβ^+^CD4^-^CD8^-^DN T cells was substantially restored by *Rictor* deletion (Fig. 1F). Finally, there were elevated, albeit to a widely variable degree, levels of multiple inflammatory cytokines in Lpr mice, such as IFNβ, IL-1β, TNFα and IL-17A, many of which were rectified in Lpr.*Rictor* T-KO mice (Fig. 1G). Therefore, the inactivation of mTORC2 in T cells significantly reverses the abnormal accumulation of T and B cells, production of immunoglobulin and inflammatory cytokine, and restores immune homeostasis in peripheral lymphoid organs of Lpr mice.

### mTORC2 in T cells promotes B cell hyper-activation

Reflecting the enhanced antibody levels, immunofluorescence staining revealed a dramatic increase of peanut agglutinin (PNA) staining in Lpr mice, suggesting increased GC reaction. The pattern of the PNA staining was variable. Some are dense aggregations like defined GC formation, but others are more diffused without defined boundaries (Fig. 2A). T cell distribution was also disorganized in Lpr lymphoid tissues. Both the increased PNA staining and disorganized T cell distribution were greatly restored by RICTOR deficiency (Fig. 2A). Flow cytometry analysis confirmed these observations. B cells from Lpr mice had highly increased expression of GC B cell marker GL-7, and significant loss of IgD expression, indicating increased B cell activation and class switch (Fig. 2B). Consistent with this notion, Lpr B cells had greatly increased IgG1 expression (Supplemental Fig. 2A). A large proportion of Lpr B cells lost expression of CD38, a NADase with strong immunosuppressive capacity^39^, and the loss of which is another characteristic of germinal center B cells (Supplemental Fig. 2A). Because plasmablasts contribute to extrafollicular response, we stained CD138, a marker of plasmablasts. The results showed a markedly increase of CD138 expression in extrafollicular region of Lpr spleens, which was reduced in Lpr.*Rictor* T-KO mice (Fig. 2C). The RICTOR-dependent increase of extrafollicular response in Lpr mice was confirmed with flow cytometry analysis. Lpr mice accumulated large number of cells with modest expression of CD138, and B220^lo^CD138^hi^ plasmablast/plasma cells, while deletion of *Rictor* in T cells dramatically suppressed the generation of these populations (Fig. 2D). Therefore, mTORC2 activity in T cells is a major contributor to B cell abnormality in Lpr mice. Taken together, our data indicate that inhibition of mTORC2 in T cells attenuates GC and plasmablast formation in Lpr mice.

**Figure 2.**
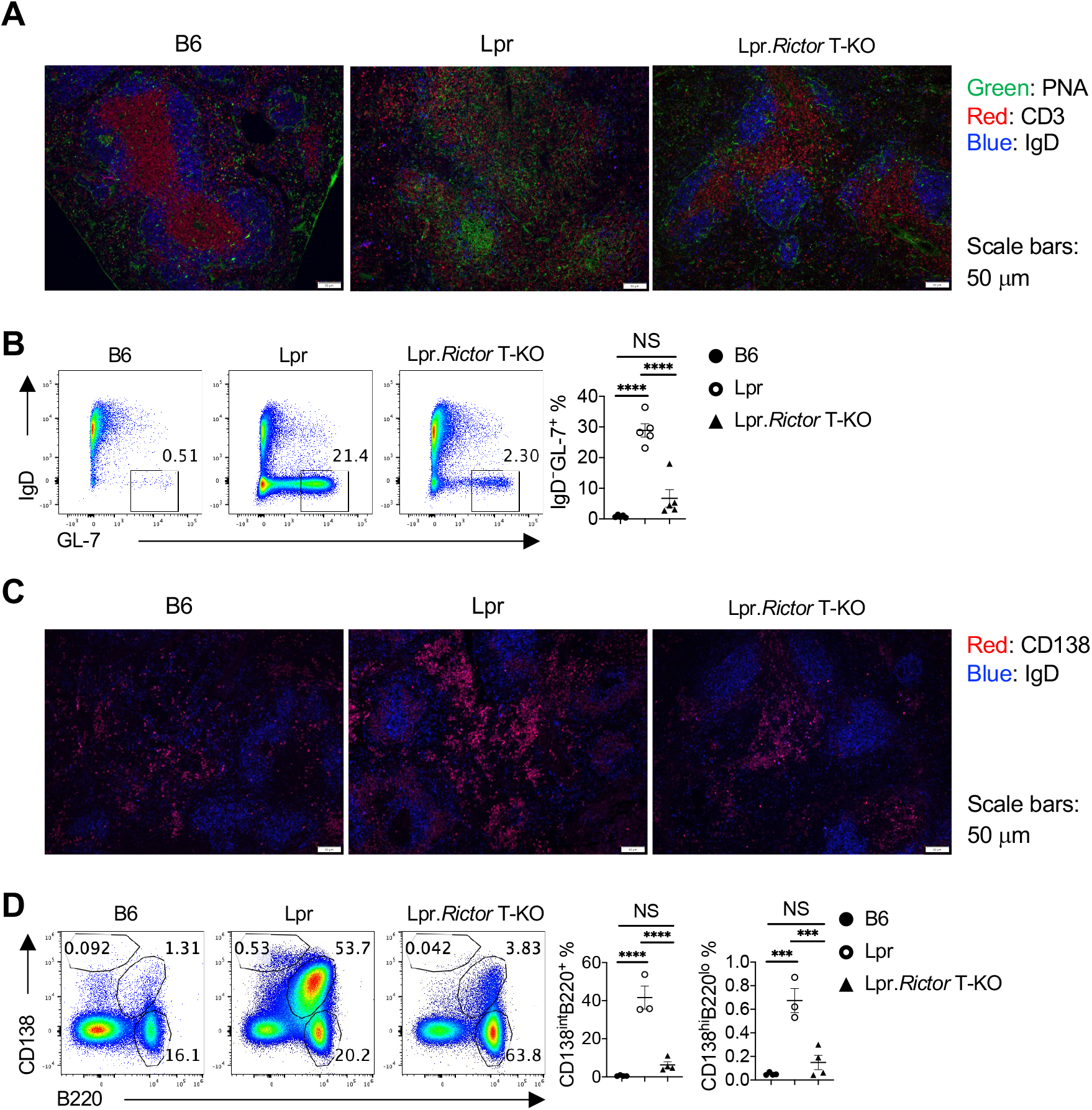
mTORC2 in T cells is required for B cell hyper-activation in Lpr mice. (A) Immunofluorescence staining of peanut agglutinin (PNA), CD3 and IgD on spleen sections from 6 months old B6, Lpr and Lpr.*Rictor* T-KO mice. (B) Flow cytometry analysis of GL-7 and IgD expression among B220^+^TCRβ^-^ cells. Right, the frequency of IgD^-^GL-7^+^ germinal center (GC) B cells. (C) Immunofluorescence staining of CD138 and IgD on spleen sections from 6 months old B6, Lpr and Lpr.*Rictor* T-KO mice. (D) Flow cytometry analysis of B220^+^CD138^int^ and B220^lo^CD138^hi^ two populations. Right, the frequencies of B220^+^CD138^int^(left) and B220^lo^CD138^hi^ (right) among total lymphocytes. (B) and (D) Cells were from pLNs of 6 months old B6, Lpr and Lpr.*Rictor* T-KO mice. NS, not significant; *** *P* < 0.001, **** *P* < 0.0001 (one-way ANOVA). Results were pooled from 3 (B and D) independent experiments. Error bars represent SEM.

### Increased Tfh differentiation in Lpr mice is dependent on mTORC2

Tfh is key for GC formation. Indeed, we observed many T cells in GC of Lpr mice from immunofluorescence staining, presumably Tfh cells (Fig. 2A). This was confirmed by flow cytometry data, which showed a strong increase of BCL6^+^CXCR5^hi^ GC Tfh and BCL6^-^CXCR5^+^ pre-Tfh cells in Lpr mice, that was significantly dependent on RICTOR expression (Fig. 3A and Supplemental Fig. 3A for gating strategy). A RICTOR-dependent CXCR5 expression was also found on TCRβ^+^CD4^-^CD8^-^ DN T cells (data not shown). Similar observations were found when using other conventional Tfh markers, PD-1 and CXCR5 (Supplemental Fig. 3B). RICTOR deficiency also reduced PSGL-1^-^CXCR5^-^ EF Tfh cell frequency (Fig. 3B), consistent with the reduction of plasmablast/plasma cells. These results indicated a critical requirement of mTORC2 signaling for autoreactive GC and EF Tfh differentiation in Lpr lupus-prone model. Similar rescue effects were observed at different ages (at 3 and 9 months, data not shown). Like SLE patients^14^, Lpr mice had highly increased ICOS expression, both on CXCR5^+^ and CXCR5^-^ T cells. Interestingly, deletion of *Rictor* failed to restore ICOS expression (measured by both positive percentages and mean fluorescence intensity), indicating that the reduced Tfh differentiation in Lpr.*Rictor* T-KO mice is not a consequence of reduced ICOS expression (Fig. 3C).

**Figure 3.**
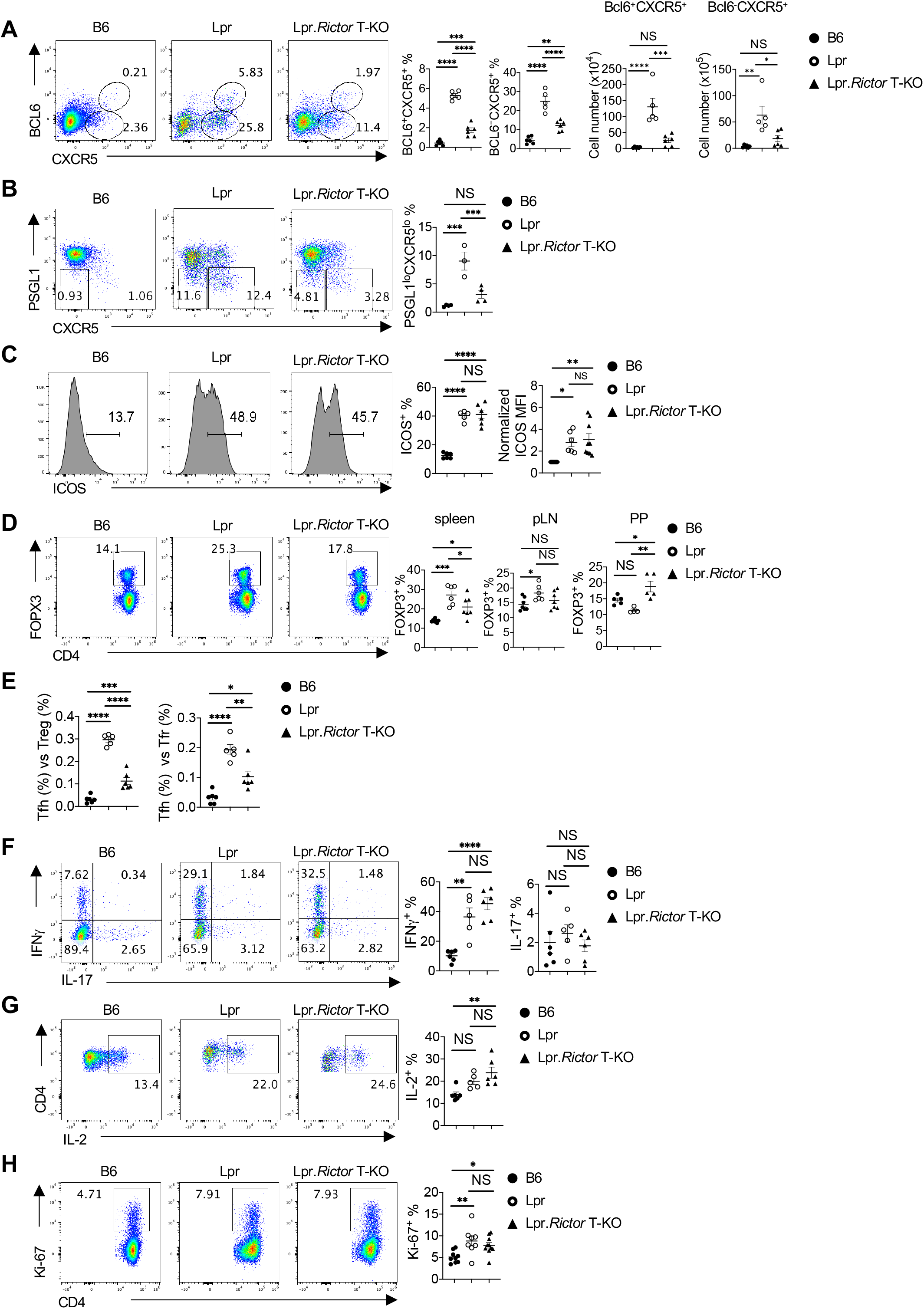
*Rictor* deletion reduces Tfh differentiation, normalizes Treg frequencies, but does not affect Th1 and Th17 differentiation or T cell proliferation. (A) Expression of CXCR5 and BCL6 on B220^-^CD4^+^ T cells. Right, the percentages (left) and absolute numbers (right) of BCL6^+^CXCR5^+^ and BCL6^-^CXCR5^+^ cells. (B) Expression of PSGL1 and CXCR5 on B220^-^ CD4^+^ T cells. Right, the summary of PSGL1^lo^CXCR5^lo^ extrafollicular (EF) Tfh cell percentages. (C) Expression of ICOS on B220^-^CD4^+^ T cells. Right, the summary of ICOS^+^ cell frequency (left) and normalized ICOS median fluorescence intensity (MFI) (right). (D) Flow cytometry analysis of Treg cells (B220^-^CD4^+^FOXP3^++^). Frequencies of FOXP3^+^ cells in CD4^+^ T cells from spleen (left), pLN (middle) and Peyer’s patches (PP) (right). (E) The ratio between Tfh and Treg frequency (left) and the ratio between Tfh and Tfr frequency (right) in CD4^+^ T cells from B6, Lpr and Lpr.*Rictor* T-KO mice. (F) Expression of IFNg and IL-17 in CD4^+^ T cells activated by PMA and ionomycin. Right, frequencies of IFNg^+^ cells and IL-17^+^ cells. (G) Expression of IL-2 in CD4^+^ T cells activated by PMA and ionomycin. Right, the frequency of IL-2^+^ cell. (H) Expression of Ki-67 in CD4^+^ T cells from B6, Lpr and Lpr.*Rictor* T-KO mice. Right, the summary of Ki-67^+^ cell frequency. (A-C, and E-H) Cells were from pLNs of 6 months old B6, Lpr and Lpr.*Rictor* T-KO mice. NS, not significant; * *P* < 0.05, ** *P* < 0.01, *** *P* < 0.001, **** *P* < 0.0001 (one-way ANOVA). Results were pooled from at least 3 (A-H) independent experiments. Error bars represent SEM.

Treg cells are key for maintaining immune tolerance and preventing autoimmunity. Counterintuitively, in an inflammatory condition, Treg percentage sometimes paradoxically increases. We found that Lpr mice had variable Treg perturbations dependent on anatomic location: while Treg percentages increased the most in spleens and modestly in peripheral lymph nodes (pLN), they trended lower in the gut mucosal site relative to WT B6 mice (Fig. 3D), in a manner consistent with the graded mTORC2 activity, gradually increasing from spleen to pLN, and to gut mucosal site^24^. In all cases, deletion of *Rictor* significantly restored the Treg homeostasis in Lpr mice (Fig. 3D). The magnitude of changes of Treg frequencies in Lpr mice were much smaller than that of Tfh cells, thus the ratio between Tfh and Treg was drastically increased in Lpr mice, suggesting an imbalance between Tfh and Tregs in Lpr mice. Such imbalance was significantly restored by mTORC2 deficiency (Fig. 3E). FOXP3^+^CXCR5^+^ follicular regulatory T (Tfr) cells are specialized to suppress Tfh cells and GC formation^40^. RICTOR deficiency alone does not impact Tfr differentiation^24^. Lpr mice had a highly increased ratio between Tfh and Tfr cells, which was partly restored in Lpr.*Rictor* T-KO mice (Fig. 3E). However, Lpr mice also had significantly increased frequency of FOXP3^+^CXCR5^+^ Tfr cells compared to B6 control mice, which was also reversed by *Rictor* deletion (Supplemental Fig. 3C). Therefore, the data indicate that RICTOR deficiency impacts Tfh/Tfr ratio primarily by reducing Tfh, not increasing Tfr.

Furthermore, we found that *Rictor* deficiency did not restore the increased IFNg expression (Fig. 3F) or alter IL-2 and IL-17 expression in CD4^+^ T cells from Lpr mice (Fig. 3F, 3G), or change the expression of lineage transcription factor Tbet and Rorgt (Supplemental Fig. 3D), consistent with our hypothesis that mTORC2 inactivation preferentially affects Tfh differentiation, but not other effector T cell lineages. These data also suggested that Tregs from Lpr.*Rictor* T-KO mice were not able to control the excessive cytokine production in Lpr CD4^+^ T cells, despite normalization of their homeostasis. Finally, the restoration of T cell phenotypes in Lpr.*Rictor* T-KO mice was likely independent of cell proliferation, because CD4^+^ T cells from Lpr.*Rictor* T-KO mice had similarly increased Ki-67^+^ percentage at steady state as those from Lpr mice (Fig. 3H). Therefore, our data showed that mTORC2 contributes to the increased GC and EF Tfh cells in Lpr mice, independent of ICOS receptor expression and T cell proliferation.

### mTORC2 inactivation in Tregs fails to restore immune homeostasis in Lpr mice

The two best described functions of mTORC2 in T cells are promoting Tfh differentiation^24,41^ and suppression of Treg function^25,29^. Thus, it is possible the above-described phenotypic improvements in Lpr.*Rictor* T-KO mice (deletion in all T cells) can be partly attributed to enhanced Treg function. To dissect whether mTORC2 contributes to systemic autoimmunity mainly through inhibiting Tregs, promoting Tfh, or both, we generated C57BL/6.*Fas^Lpr^Foxp3*^Cre^*Rictor*^fl/fl^ (Lpr.*Rictor* Treg-KO) mice, in which *Rictor* was specifically deleted in Treg cells (Supplemental Fig. 4A). Although *Rictor* deficiency in Tregs can boost Treg suppressive function in the absence of mTORC1^25^, *Pten*^29^ or *Foxp3*^31^, it also reduces peripheral Treg frequency compared to WT controls^25^. Lpr.*Rictor* Treg-KO mice also had reduced Treg percentage compared to Lpr mice, although their Treg percentage was similar to Lpr.*Rictor* T-KO mice (Fig. 4A). We compared the immunological phenotypes between Lpr, Lpr.*Rictor* T-KO, and Lpr.Rictor Treg-KO mice. To our surprise, Lpr.*Rictor* Treg-KO mice had equivalent lymphoplasia found in Lpr mice, as illustrated by the lymphadenopathy (Fig. 4B). The expansion of B220^+^TCRβ^+^ cells was not affected by Treg specific deletion of *Rictor* (Fig. 4C). Similarly, the elevated Tfh, pre-Tfh and GC B cells in Lpr.*Rictor* Treg-KO mice were all at a comparable level as those found in Lpr mice, in sharp contrast to Lpr.*Rictor* T-KO mice (Fig. 4D, E). Because of the elevated Tfh frequency, Lpr.*Rictor* Treg-KO mice retained a high Tfh vs Treg ratio similar to that of Lpr mice (Fig. 4F). Quantification of immunoglobulin concentrations revealed that Lpr.*Rictor* Treg-KO mice had highly elevated antibody levels, some of which (including IgG1 and IgG2a) even surpassed those in Lpr mice, whereas Lpr.*Rictor* T-KO mice had significantly reduced IgG2a and IgG2b levels and IgG1 level trending lower (but not significantly) compared to those in Lpr mice (Fig. 4G). These data support the idea that mTORC2-dependent overactivation of Tfh, but not mTORC2-mediated impairment of Treg cells, contributes to the systemic lymphoproliferation in Lpr mice.

**Figure 4.**
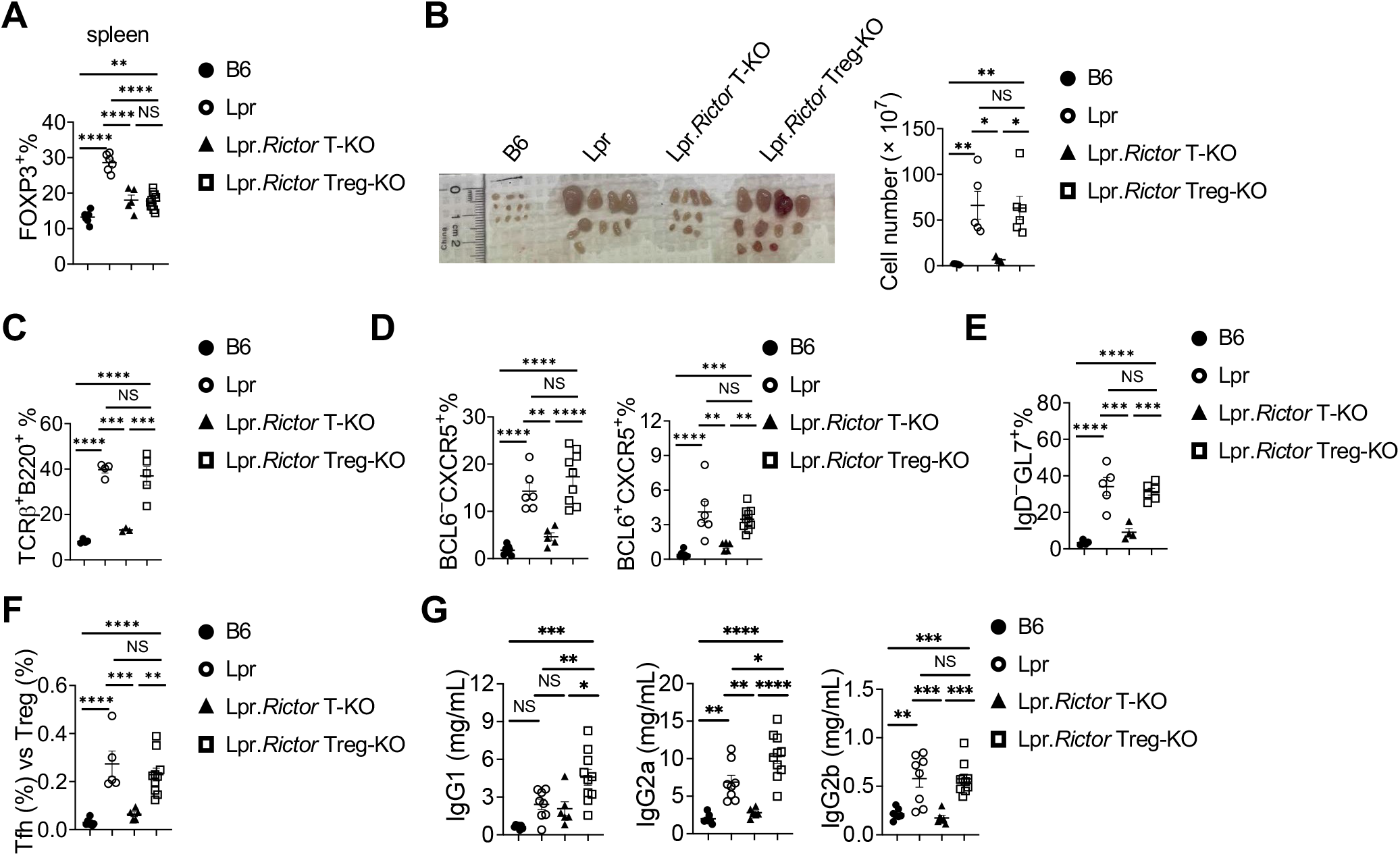
*Rictor* deletion in Treg is unable to restore immune dysregulation in Lpr mice. (A) Frequency of FOXP3^+^ Treg cells in splenic CD4^+^ T cells. (B) Image of peripheral lymph nodes taken from 6 months old B6, Lpr, Lpr.*Rictor* T-KO mice and Lpr.*Rictor* Treg-KO. Right, summary of total cellularity of lymph nodes. (C) Frequency of B220^+^TCRβ^+^ cells in pLN among B6, Lpr, Lpr.*Rictor* T-KO mice and Lpr.*Rictor* Treg-KO. (D) Frequencies of pre-Tfh (BCL6^-^ CXCR5^+^) and Tfh (BCL6 ^+^CXCR5^+^) cells in pLN. (E) Frequency of GC B cells. (F) The ratio between Tfh and Treg frequency in CD4^+^ T cells from B6, Lpr, Lpr.R*ictor* T-KO and Lpr.*Rictor* Treg-KO mice. (G) Serum concentrations of immunoglobulin IgG1, IgG2a, and IgG2b were measured by LEGENDplex. NS, not significant; * P < 0.05, ** P < 0.01, *** P < 0.001, **** P < 0.0001 (one-way ANOVA). Results were pooled from at least 3 (A-G) independent experiments. Error bars represent SEM.

### Inactivation of mTORC2 restores T cell glucose metabolism

Recent studies indicate that lupus development is associated with dysregulation of T cell immunometabolism^42^. mTORC2 is known as one of the key regulators for cell metabolism. We compared the basal respiration and glycolysis of T cells among B6, Lpr and Lpr.*Rictor* T-KO mice at different ages under TCR signaling stimulation. Consistent with recent publications^43,44^, T cells from 2 months old Lpr mice had modestly increased basal respiration and glycolysis (Fig. 5A, 5B). Strikingly, T cells from 3 and 9 months old Lpr mice had dramatically decreased basal respiration and glycolysis, relative to B6 mice (Fig. 5A, 5B). The opposite metabolic activities between 2 and 3 months old Lpr mice T cells were associated with drastic changes of naive and effector T cell composition at different ages of Lpr mice during lupus development (Supplemental Fig. 5A). Lpr.*Rictor* T-KO mice had much lower levels of basal respiration and glycolysis compared to B6 mice at all ages (Fig. 5A, 5B). The reduced metabolic activity in primary culture of T cells from older Lpr and Lpr. *Rictor* T-KO mice (compared to 2 months old mice) was consistent with reduced T cell proliferation during primary culture (Supplemental Fig. 5B). Because anergic T cells are known to have stunted metabolic activity and reduced glycolysis^45,46^, we tested whether the reduced metabolism in 6 months old Lpr T cells was associated with increased anergic T cells. Indeed, we found drastically increased anergy (FR4^+^CD73^+^)^47^ markers on Lpr CD4^+^ T cells at steady state, which was mildly reduced in Lpr.*Rictor* T-KO mice (Supplemental Fig. 5C). Costimulatory signaling from ICOS is important to limit T cell anergy^48^. Furthermore, we have previously shown that mTORC2 functioned as downstream of ICOS receptor to promote Tfh differentiation, which could be modeled by restimulating previously activated T cells with anti-CD3/anti-ICOS^24^. Interestingly, Lpr T cells had enhanced proliferation during secondary activation with anti-CD3/anti-ICOS regardless of the ages (Supplemental Fig. 5D, and data not shown), suggesting that re-stimulation from TCR and ICOS signaling not only reversed the proliferation defect, and possibly broke the anergic status, in Lpr T cells, but also promote more accumulation of Lpr T cells relative to WT control. All these defects were not markedly affected by RICTOR deficiency. mTORC2 promotes TCR/ICOS mediated glucose metabolism during Tfh differentiation^24^. Therefore, we examined the T cell glycolysis after anti-CD3/anti-ICOS secondary stimulation. Lpr CD4^+^ T cells exhibited a significant increase of compensatory glycolytic rate relative to B6 controls, regardless of the ages. *Rictor* deletion substantially reduced both basal and compensatory glycolytic rate in Lpr CD4^+^ T cells (Fig. 5C). Therefore, mTORC2 contributes to the TCR/ICOS mediated enhanced glycolysis in Lpr T cells without affecting T cell proliferation.

**Figure 5.**
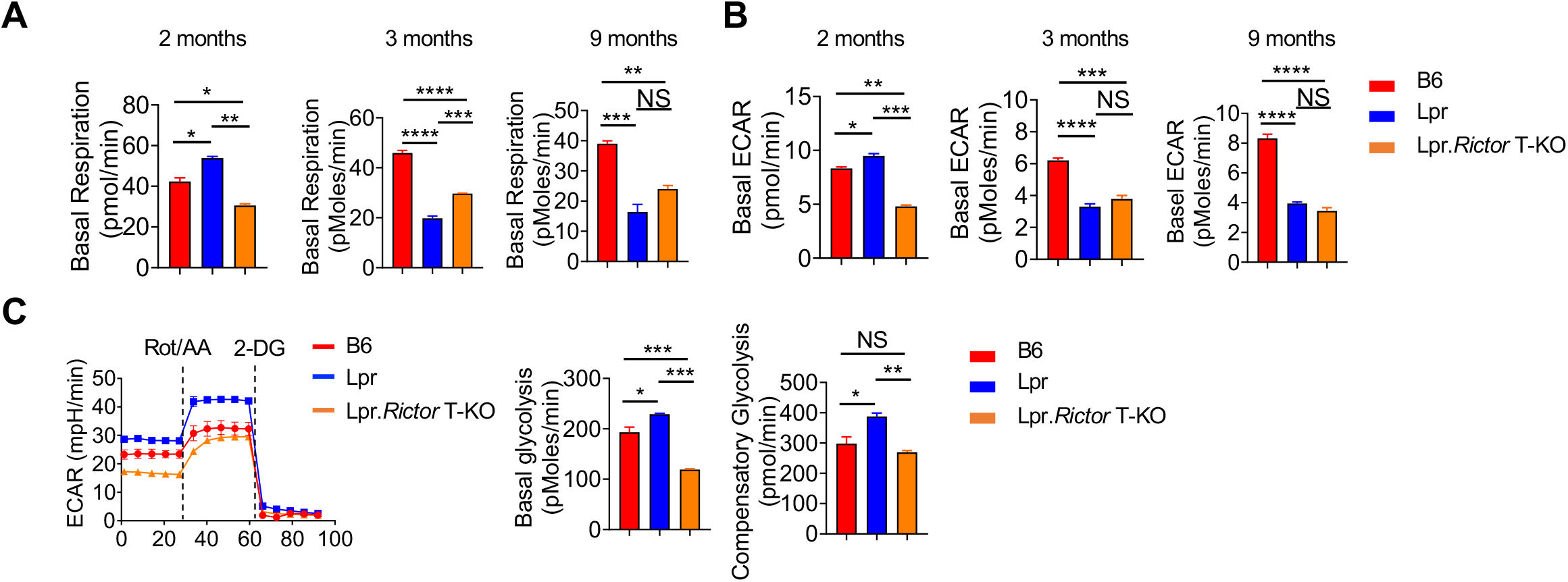
mTORC2 supports TCR/ICOS mediated glucose metabolism in Lpr T cells. (A) The basal respiration and (B) basal extracellular acidification rate (ECAR) of CD4^+^ T from 2-, 3-, and 9-months old mice cells under anti-CD3/anti-CD28 overnight activation, respectively. (C) The ECAR of CD4^+^ T cells followed by sequential anti-CD3/anti-CD28, and anti-CD3/anti-ICOS stimulation measured by glycolytic rate assay. Cells were from pLNs of 6 months old B6, Lpr and Lpr.*Rictor* T-KO mice. Right, summaries of basal glycolysis and compensatory glycolysis. Rot, Rotenone; AA, Antimycin A; 2-DG, 2-deoxyglucose. NS, not significant; * *P* < 0.05, ** *P* < 0.01, *** *P* < 0.001, **** *P* < 0.0001 (one-way ANOVA). Results were presentative of 2 (A-B) and 3 (C) independent experiments. Error bars represent SEM.

### mTORC2 functions downstream of type I IFN to promote CD69 expression and suppress T cell egress

Previous studies have identified that type I IFN contributes to the lupus phenotypes in Lpr mice^49,50^. We also observed increased type I IFN in Lpr mouse serum (Fig. 1G). Interestingly, we found that addition of IFNa significantly enhanced anti-CD3/anti-CD28 induced mTORC2 target p-AKT_473_, but not mTORC1 target p-S6, at 6 h stimulation, which was completely dependent on RICTOR (Fig. 6A). IFNa alone could also induced weak p-AKT_473_, but the combination of anti-CD3/anti-CD28 and IFNa appeared to be the most potent activators of RICTOR dependent p-AKT_473_ (Fig. 6A, 6B). RICTOR deficiency did not markedly affect canonical IFNα–STAT1 signaling or mTORC1 activation (Fig. 6A, Supplementary Fig. 6A). Similar results were observed using IFNβ (Supplementary Fig. 6B). Thus, type I IFN, in synergy with anti-CD3/anti-CD28, promoted p-AKT_473_ in a RICTOR dependent manner. We also observed similar synergy between TCR and type I IFN on mTORC2 activation in human CD4^+^ T cells (Fig. 6C). Type I IFN is known to suppress lymphocyte egress from lymph nodes to blood, partly through induction of CD69 expression^7,8^. Poly(I:C) is a potent inducer of type I IFN. It induces CD4^+^ T cell lymphopenia in an IFNAR dependent manner^8^. Indeed, administering poly(I:C) induced potent CD69 expression on blood CD4^+^ T cell (Fig. 6D), and induced CD4^+^ T cell lymphopenia (Fig. 6E). Both phenotypes were significantly reversed in CD4^+^ T cells from *Cd4^Cre^Rictor*^fl/fl^ mice. Such phenotype was not observed in CD19^+^ B cells, in which RICTOR was intact (Supplemental Fig. 6C), indicating that type I IFN promotes CD4^+^ T cell lymphopenia partly through T cell intrinsic mTORC2. Consistent with these observations, Lpr CD4^+^ T cells had increased CD69 level, which was restored in Lpr.*Rictor* T-KO mice in both lymph nodes and blood (Fig. 6F, Supplemental Fig. 6D). Importantly, the severe CD4^+^ T cell lymphopenia phenotype in Lpr mice was also modestly, but significantly, reversed by RICTOR deficiency (Fig. 6G, Supplemental Fig. 6E). The rescue effect on CD69 expression was consistently more pronounced than that on CD4^+^ T cell lymphopenia (Fig. 6D-G), suggesting that a large part of lymphopenia phenotype in Lpr mice may be CD69 independent. Altogether, CD4^+^ T cells from Lpr mice exhibited increased mTORC2 activity. Type I IFN, together with TCR signaling, activates mTORC2 to induce CD69 expression on CD4^+^ T cells and blocks CD4^+^ T cell egress into blood. RICTOR deficiency partially restores CD4^+^ T cell lymphopenia in Lpr mice.

**Figure 6.**
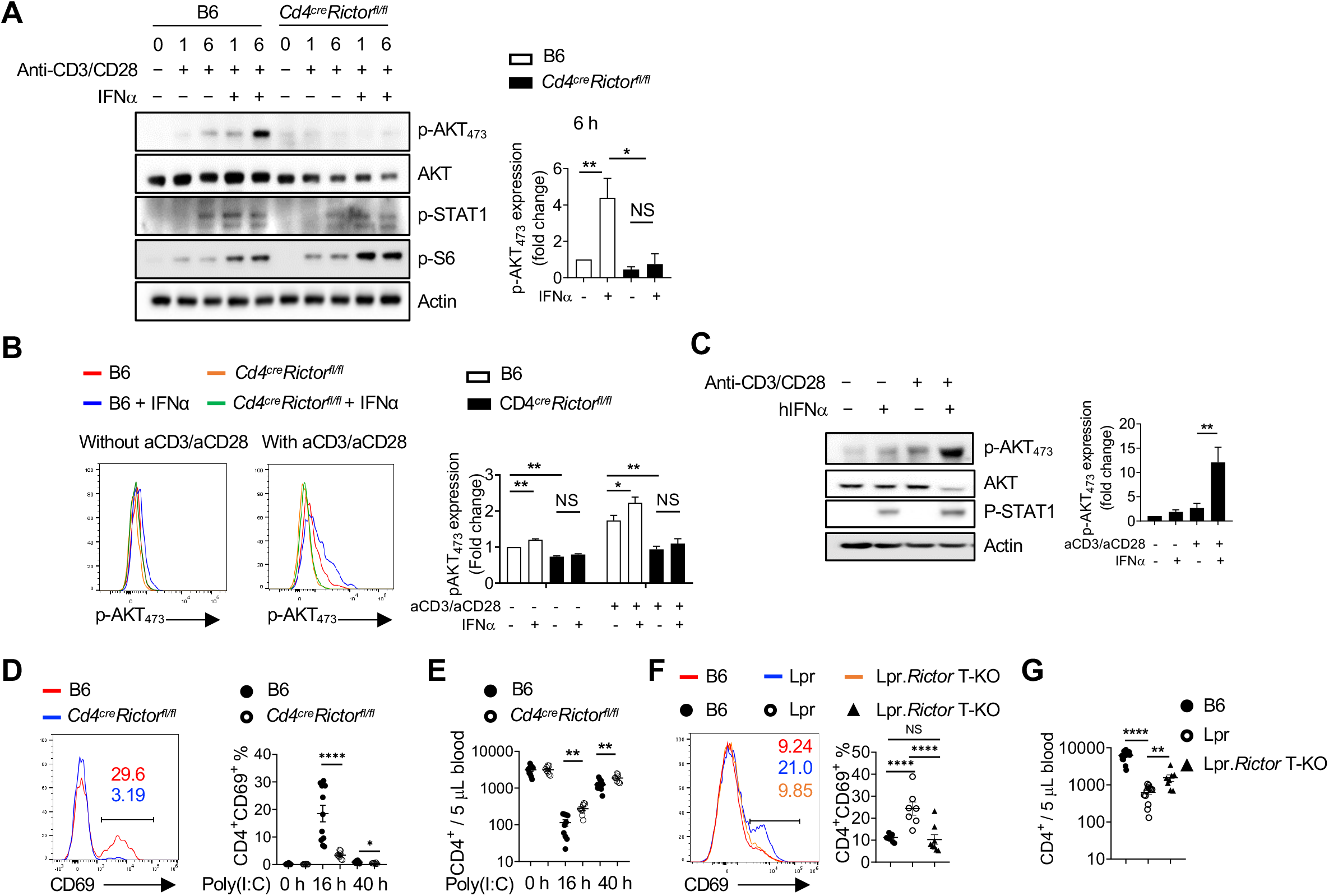
Type I IFN synergizes with TCR to promote CD69 expression and suppress T cell egress. (A) Immunoblot analysis of p-AKT_473_, AKT (pan), p-STAT1 and p-S6 in CD4^+^ T cells from B6 and *Cd4*^cre^*Rictor*^fl/fl^ mice stimulated with or without anti-CD3/anti-CD28 in the presence or absence of IFNα for 1 and 6 hours. Right, summaries of the relative p-AKT_473_ expression (normalized to total AKT and then compared with baseline B6 CD4^+^ T cells) stimulated for 6 hours. (B) Flow cytometry analysis of p-AKT_473_ expression in CD4^+^ T cells treated with IFNα alone, or in combination with anti-CD3/anti-CD28 overnight. Right, summary of the relative pAKT_473_ expression (normalized to that in B6 CD4^+^ T cells without any stimulation). (C) Immunoblot analysis of p-AKT_473_, AKT (pan), and p-STAT1 in human CD4^+^ T cells with or without anti-CD3/anti-CD28 activation in presence or absence of human IFNα for 3 hours. Right, summary of the relative p-AKT_473_ expression normalized to total AKT and then compared with baseline. (D) and (E) B6 and *Cd4^cre^Rictor*^fl/fl^ mice were administered with poly(I:C) intraperitoneally. (D) Expression of CD69 in blood CD4^+^ T cells from B6 and *Cd4^cre^Rictor*^fl/fl^ mice after poly(I:C) administration. Numbers indicate the percentages of CD69^+^CD4^+^ T cells. Right, summary of CD69^+^ percentage in CD4^+^ T cells at baseline or treated with poly(I:C) for 16 h and 40 h. (E) Blood CD4^+^ T cell counts were determined before and after poly(I:C) treatment. (F) Expression of CD69 in CD4^+^ T cells from B6, Lpr and Lpr.*Rictor* T-KO mice. Right, summary of CD69^+^ percentages among pLN CD4^+^ T cells. (G) Blood CD4^+^ T cell counts were determined in 4-6 months old B6, Lpr and Lpr.*Rictor* T-KO mice. NS, not significant; * *P* < 0.05, ** *P* < 0.01, *** *P* < 0.001, **** *P* < 0.0001 (A, B, D and E, unpaired Student’s *t* test, C, F and G, one-way ANOVA). Results were presentative of 4 (A, C), or pooled from at least 3 (A-G) independent experiments. Error bars represent SEM.

## DISCUSSION

In this study, we experimentally tested the hypothesis that genetic targeting mTORC2 in T cells may benefit systemic autoimmunity using a classic systemic lymphoproliferative mouse model, Lpr mice. We made several key observations: 1) T cells in Lpr mice have elevated mTORC2 activity and mTORC2 inhibition through genetic deletion of RICTOR in T cells can significantly ameliorate immunopathology of Lpr mice, including increased GC B cells and extrafollicular B cells; 2) RICTOR deficiency restores Tfh and Treg balance in Lpr mice, without significant alteration of T cell proliferation and Th1 and Th17 lineages; 3) RICTOR deficiency in Tregs fails to rectify most of the lymphoproliferative phenotypes in Lpr mice; 4) RICTOR deficiency reduces ICOS-mediated glycolysis in Lpr T cells. 5) Type I IFN promotes mTORC2 activation in CD4^+^ T cells partly to drive CD69 expression and T cell lymphopenia. Thus, our study suggests that mTORC2 might be a potential therapeutic target for systemic autoimmunity.

Our genetic studies demonstrated that mTORC2-dependent overactivation of Tfh cells, but not impairment of Tregs, contributes to systemic autoimmunity in Lpr mice. Although previous study showed deletion of *Rictor* in Tregs restores Treg suppressive activity, including the Tfh suppressing ability, our data demonstrated that deletion of mTORC2 in Tregs does not improve their ability to suppress Tfh cells in Lpr mice. These observations suggest that the ability to promote Tfh differentiation is likely a more dominant function of mTORC2 in systemic autoimmunity, at least in Lpr background. Furthermore, it is worth noting that while RICTOR deficiency greatly reduced Tfh differentiation in Lpr mice, it did not significantly affect Th1 and Th17 subsets, T cell IL-2 production and proliferation. These observations are consistent with our hypothesis that mTORC2 specifically promotes Tfh differentiation without substantial effects on T cell activation, proliferation, and other effector T cell lineages. This contrasts with mTORC1, which is required for T cell quiescence exit and all effector T cell lineages^22^. Thus, our data suggest that targeting mTORC2 may specifically suppress Tfh lineage, which may have relatively smaller immunosuppressive side effects, compared to inhibition of mTORC1.

It has long been established that exuberant type I IFN activity contributes to lupus development. Yet, the molecular mechanism through which type I IFN impact T cells in lupus remains largely unknown. We show that type I IFN may be one of the upstream signals that activate mTORC2 in T cells. One of the downstream effects of type I IFN-mTORC2 activation is to promote CD69 expression and T cell lymphopenia. However, our data also showed that mTORC2 is only part of type I IFN program that controls T cell egress, because RICTOR deficiency had a relatively small rescue effect on poly(I:C) induced lymphopenia or lymphopenia in Lpr mice. The prevailing theory on the pathogenesis of the lymphopenia in lupus has been centered on antilymphocyte antibodies^51^. But a large-scale multiplex single cell RNA-seq analysis demonstrated a clear inverse correlation between type I IFN activity and circulating naïve CD4^+^ T cells in SLE patients^12^. Our study corroborates with such observation indicating that elevated type I IFN production may be another contributing factor for the lymphopenia phenotype in lupus partly by potentiating mTORC2 activation. Consistent with this idea, a recent clinical trial showed that type I IFN receptor blockade can correct the T cell lymphopenia phenotype in patients with SLE^52^. One major limitation of our study is that our conclusions are based on one genetic mouse model, Lpr mice. Lpr mice carry spontaneous mutation in *Fas* gene. On C57BL/6 background, *Fas* mutation does not lead to full-blown clinical symptoms characteristic of human SLE, although many of the immune cell hyperactivation phenotypes resemble SLE. Further investigations using other mouse models and patient samples are needed. Altogether, our results support the notion that mTORC2 could be a novel target for systemic autoimmunity.

## ACKNOWLEDGEMENTS

We thank Drs. Virginia Shapiro and Jie Sun for thoughtful discussions. This work was partly supported by NIH R01AR077518 and R01Al162678, Mayo Foundation for Medical Education and Research, and Center for Biomedical Discovery at Mayo Clinic, and Lupus Research Alliance (696599, to H.Z.).

## Author contributions

H.Z. conceived and designed the study. X.Z. designed the experiments and performed most of the *in vitro* and *in vivo* experiments. H.Q. and M.L. performed ELISA analyses and cellularity calculations. Y.L. performed immunofluorescence assays on spleen samples. X.Z. performed some of the immunoblot assays. S.A. and M.A. analyzed histological samples. C.D. and A.D. performed immunofluorescence assays on kidney samples.

## Abbreviations used in this article

Tfh: follicular helper T cells
Treg: regulatory T cells
SLE: systemic lupus erythematosus
mTOR: mechanistic target of rapamycin
IFN: interferon
GC: germinal center
EF: extrafollicular
Tfr: follicular regulatory T cells
ICOS: inducible T-cell costimulator

---

**Supplemental Figure 1.**
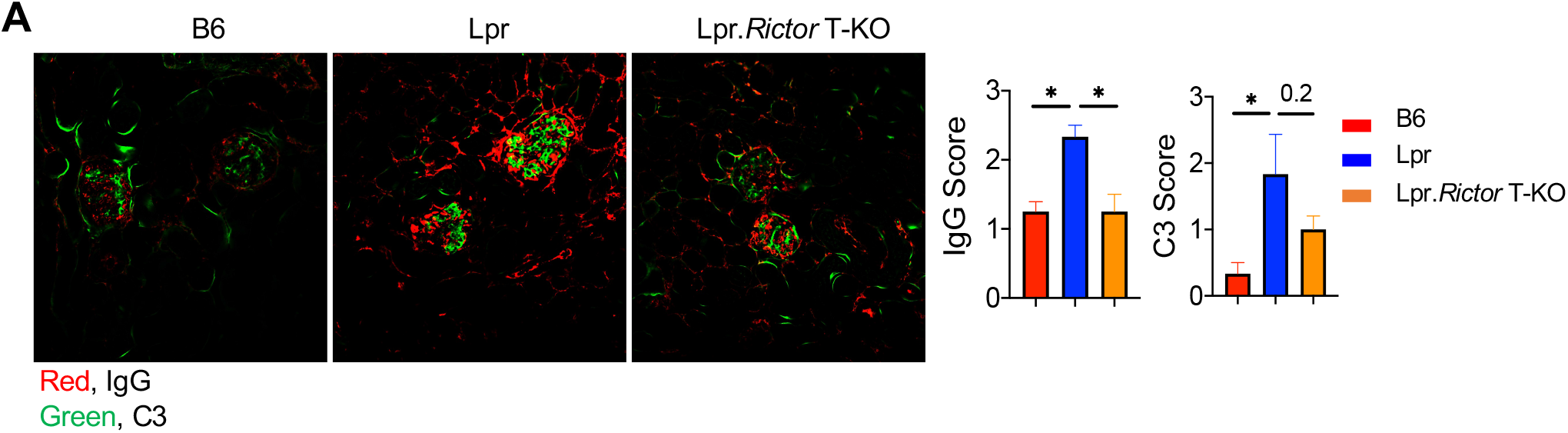
(A) Immunofluorescence staining of IgG (Red) and C3 (Green) on mice kidney sections. Right, the scores of IgG and C3 deposition. * *P* < 0.05 (one-way ANOVA). Results were pooled from 2 independent experiments. Error bars represent SEM.

**Supplemental Figure 2.**
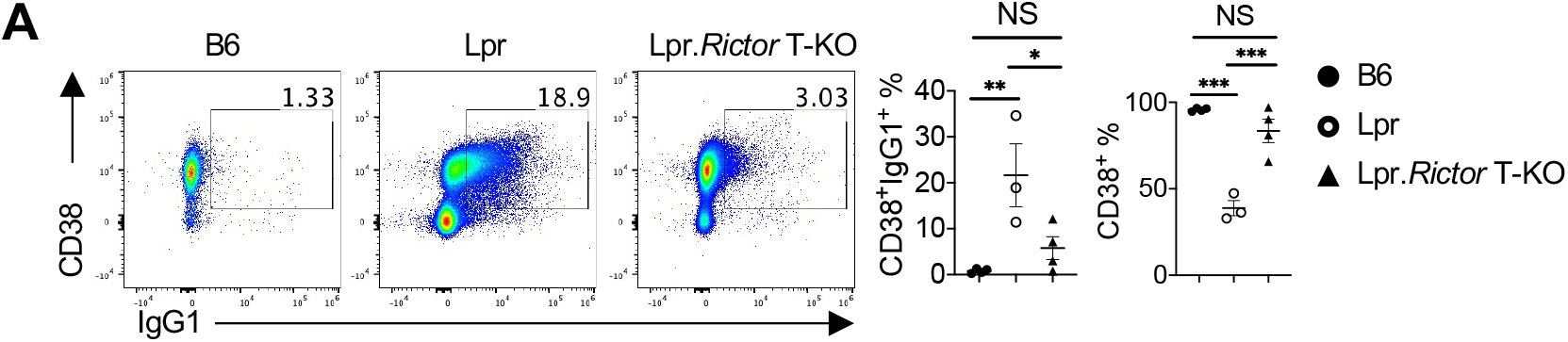
(A) Expression of CD38 and IgG1 expression on B220^+^TCRb^-^ B cells. Right, the frequencies of CD38^+^IgG1^+^ B cells (left) and CD38^+^ percentages (right). (NS, not significant; * *P* < 0.05, ** *P* < 0.01, *** *P* < 0.001, **** *P* < 0.0001, one-way ANOVA, Results were pooled from 3 independent experiments. Error bars represent SEM.

**Supplemental Figure 3.**
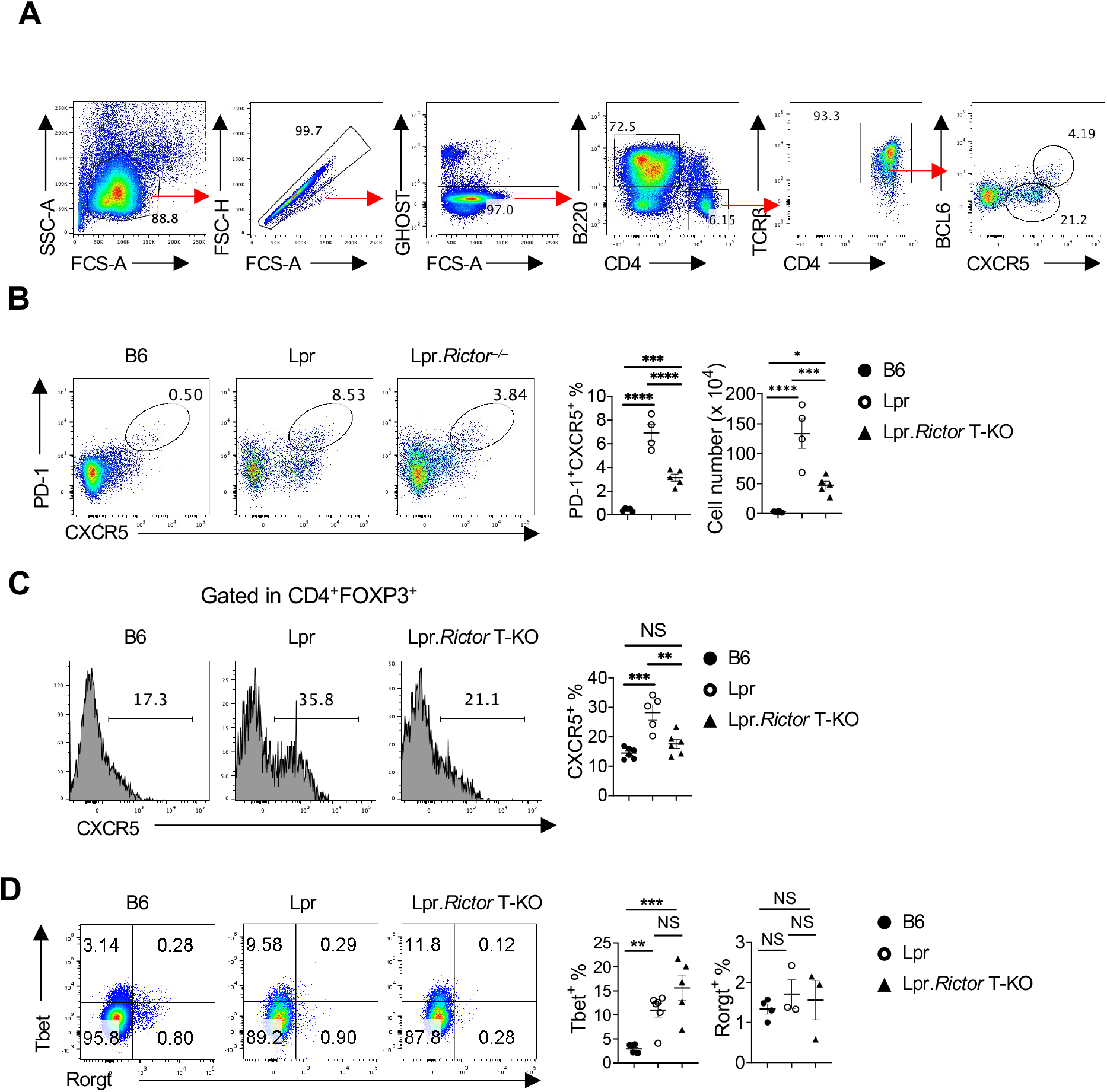
(A) Gating strategy for analysis of Tfh cells.(B) Flow cytometry analysis of CXCR5 and PD-1 on B220 ^-^CD4^+^ T cells. Right, the percentages of PD-1^+^CXCR5^+^ cells. (C) Expression of CXCR5 in Tregs. Right, summary of CXCR5^+^ Tfr cell percentages. (D) Expression of CD25 on FOXP3^+^ Tregs. Numbers indicate the percentages of CD25^-^ Tregs. Right, summary of the percentages of CD25^-^ Tregs. (F) Expression of Tbet and Rorgt on on B220^-^CD4^+^ T cells. Right, summary of Tbet^+^ and Rorgt^+^ cell percentages. Cells were from peripheral lymph nodes (pLNs) of 6 months old B6, Lpr and Lpr.*Rictor*T-KO mice. NS, not significant; ** *P* < 0.01, *** *P* < 0.001, **** *P* < 0.0001 (one-way ANOVA). Results were pooled from at least 3 independent experiments. Error bars represent SEM.

**Supplemental Figure 4.**
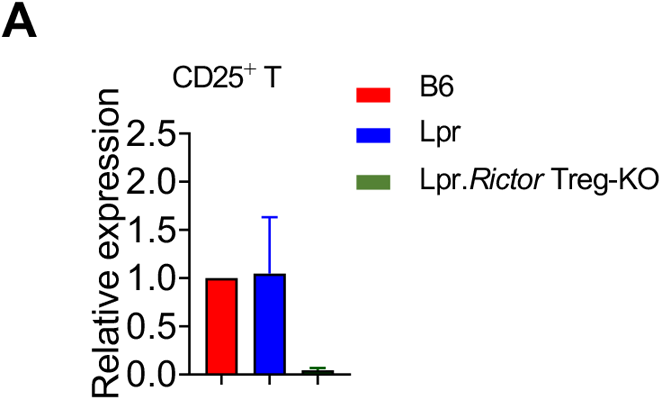
(A) Expression of *Rictor* in sorted CD25^+^ CD4^+^ T cells from B6, Lpr, and Lpr.*Rictor* Treg-KO mice. Data represent 2 independent experiments.

**Supplemental Figure 5.**
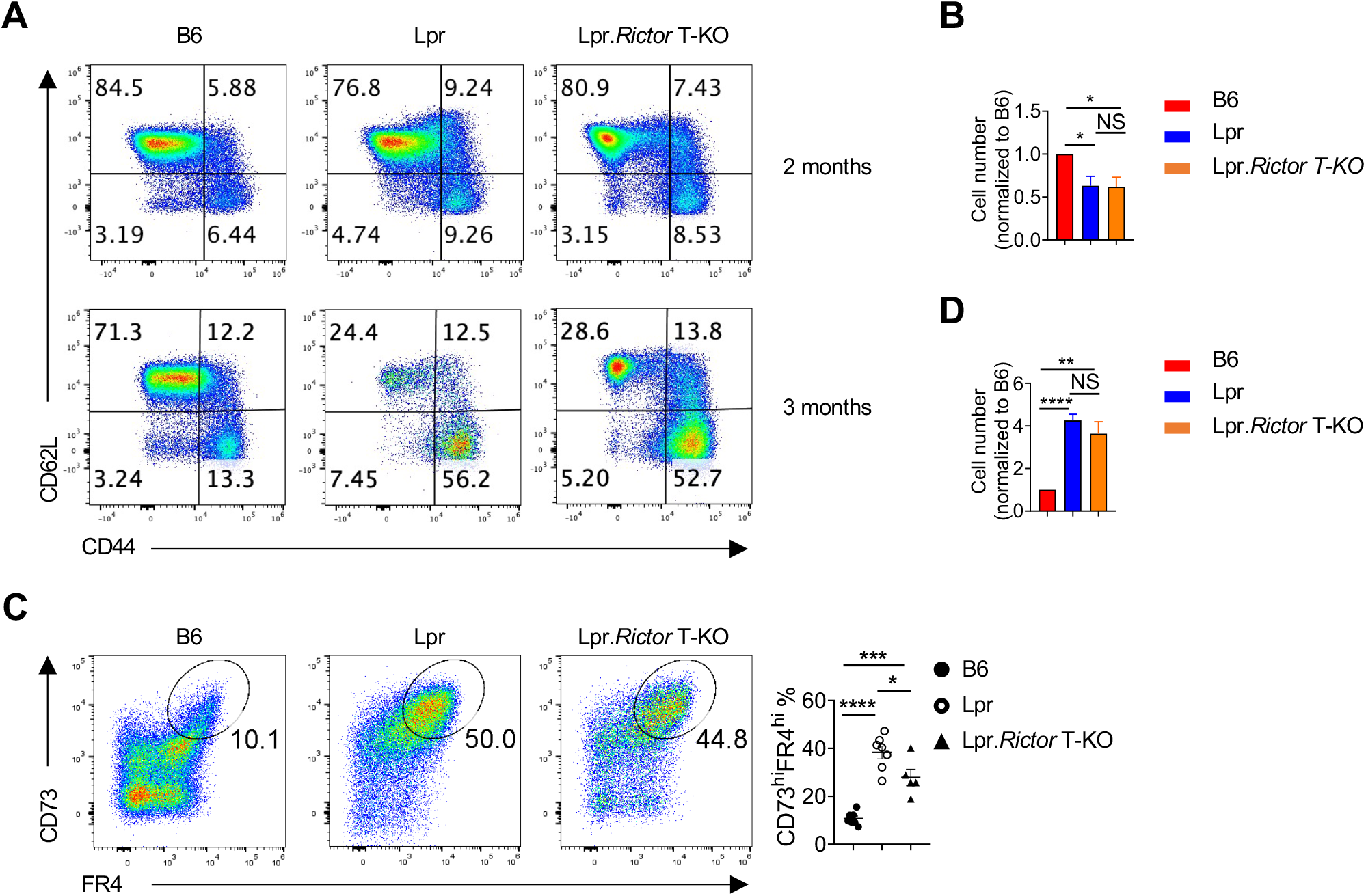
(A) Flow cytometry analysis of CD44 vs. CD62L in B220^-^CD4^+^ T cells among 2 and 3 months old B6, Lpr and Lpr.*Rictor* T-KO mice, respectively. (B) Summary of cell numbers for B220^-^ CD4^+^ T cells with anti-CD3/anti-CD28 activation for 72 hours. Numbers were normalized to B6 control. (C) Flow cytometry analysis of CD73 vs. FR4 expression in B220^-^CD4^+^ T cells among 6 months old B6, Lpr and Lpr.*Rictor* T-KO mice. Right, summaries of frequencies of CD73^hi^FR4^hi^ populations. (D) Summary of cell numbers for B220^-^CD4^+^ T cells activated with anti-CD3/anti-CD28, and then with secondary anti-CD3/ICOS stimulation. Numbers were normalized to B6 control. NS, not significant; * P < 0.05, ** P < 0.01, *** *P* < 0.001, **** P < 0.0001 (one-way ANOVA). Results were pooled from at least 3 independent experiments. Error bars represent SEM.

**Supplemental Figure 6.**
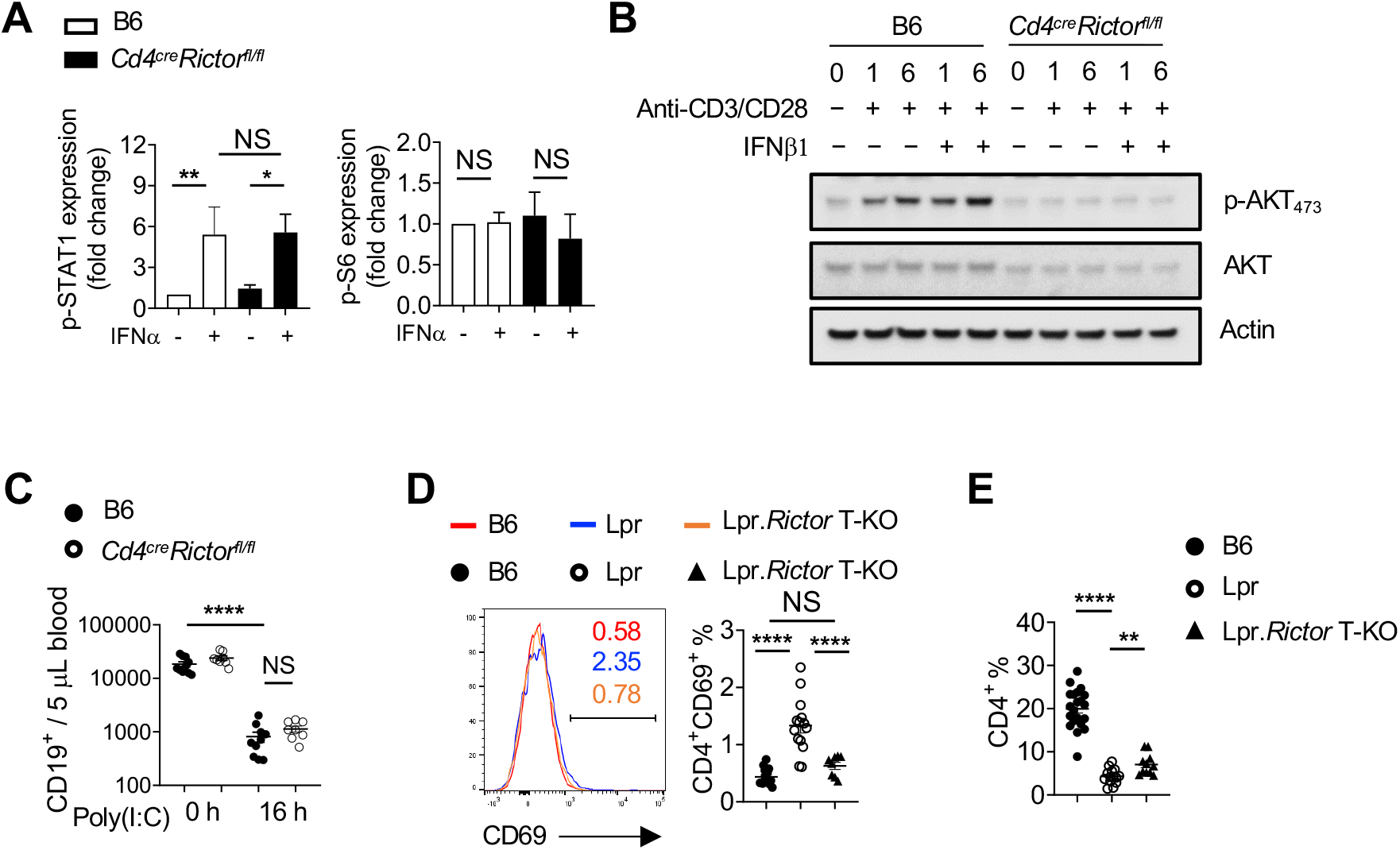
(A) Immunoblot analysis of p-STAT1 and p-S6 expression in CD4^+^ T cells from B6 and *Cd4*^cre^*Rictor*^fl/fl^ mice treated with or without IFNα in presence of anti-CD3/anti-CD28 for 6 hours. The relative expressions were normalized to them in B6 CD4^+^ T cells without any stimulation. (B) Immunoblot analysis of p-AKT_473_, p-STAT1 and Actin in CD4 T cells from B6 and *cd4*^cre^*Rictor*^fl/fl^ mice activated with anti-CD3/anti-CD28 in the presence or absence of IFNb. (C) Blood CD19^+^ B cell counts were determined before and after poly(I:C) treatment in B6 and *Cd4*^cre^*Rictot*^fl/fl^ mice. (D) Expression of CD69 was measured by flow cytometry in blood CD4^+^ T cells from 4-6 months old B6, Lpr and Lpr.*Rictor* T-KO mice. Right, summary of CD69^+^ percentages among blood CD4^+^ T cells. (E) Percentages of CD19^-^CD4^+^ T cells in blood from B6, Lpr and Lpr.*Rictor*T-KO mice among total lymphocytes. NS, not significant; * *P* < 0.05, ** *P* < 0.01, *** *P* < 0.001, **** *P* < 0.0001 (A, unpaired Student’s *t* test, C-E, one-way ANOVA). Results were pooled from at least 3 independent experiments. Error bars represent SEM.

